# Dietary Restriction Fails to Extend Lifespan of *Drosophila* Model of Werner Syndrome

**DOI:** 10.1101/2023.09.20.558691

**Authors:** Eileen Sember, Ranga Chennakesavula, Breanna Beard, Mubaraq Opoola, Dae-Sung Hwangbo

## Abstract

Werner syndrome (WS) is a rare genetic disease in humans, caused by mutations in the *WRN* gene that encodes a protein containing helicase and exonuclease domains. WS is characterized by symptoms of accelerated aging in multiple tissues and organs, involving increased risk of cancer, heart failure, and metabolic dysfunction. These conditions ultimately lead to the premature mortality of patients with WS. In this study, using the null mutant flies (*WRNexo^Δ^*) for the gene WRNexo (CG7670), homologous to the exonuclease domain of WRN in humans, we examined how diets affect the lifespan, stress resistance, and sleep/wake patterns of a *Drosophila* model of WS. We observed that dietary restriction (DR), one of the most robust non-genetic interventions to extend lifespan in animal models, failed to extend the lifespan of *WRNexo^Δ^* mutant flies and even had a detrimental effect in females. Interestingly, the mean lifespan of *WRNexo^Δ^* mutant flies was not reduced on a protein-rich diet compared to that of wild-type flies. Compared to wild type control flies, the mutant flies also exhibited altered responses to DR in their resistance to starvation and oxidative stress, as well as changes in sleep/wake patterns. These findings show that the WRN protein is necessary for mediating the effects of DR and suggest that the exonuclease domain of WRN plays an important role in metabolism in addition to its primary role in DNA repair and genome stability. Our results also raise the possibility that diet-mediated interventions could ameliorate the symptoms of WS.

## Introduction

Werner syndrome (WS; OMIM# 277700) is an autosomal recessive disease that is characterized by accelerated aging with a high incidence of cancer, heart disease, low muscle mass, and metabolic syndromes including type II diabetes, dyslipidemia, and fatty liver [1]. Recent data shows that the median life expectancy of a person with WS is 54 years old [1]. The leading causes of WS-associated death are cancer and myocardial infarction [2]. WS is caused by mutations in the WRN gene [1, 2]. The WRN protein is a member of the RecQ family of helicases and plays important roles in DNA structure and function including repair, replication, recombination, and telomere maintenance [1, 2]. In humans, the WRN protein contains domains for a 3’ to 5’ ATP-dependent DNA helicase activity and a 3’ to 5’ DNA exonuclease activity. Over 80 different types of mutations in various regions of WRN gene has been discovered in WS patients, including substitution mutations within the exonuclease domain [1–3]. To date, several animal models of WS have been generated (reviewed in [4, 5]). For example, mutant mice with only the helicase domain deleted (*Wrn*^Δ*hel*^/^Δ*hel*^) and null mutant mice with both the helicase and exonuclease activities abolished (*Wrn^-/-^*) exhibited a mean lifespan reduction ranging from 10% to 16.5% compared to wild-type (WT) mice [6–8]. These observations suggest that both domains are required for normal lifespan in mice and contribute to premature mortality when not fully functional. Unlike mice, the helicase and exonuclease domains of the human WRN protein are encoded by independent genes in worms and flies [9, 10]. Loss-of-function mutant worms for *wrn-1*, the *C. elegans* ortholog of the helicase domain of the human *WRN*, also showed a strong reduction in lifespan [11].

Although multiple animal studies have reported roles of the helicase domain of WRN protein in accelerated aging phenotypes and shortened lifespan, relatively less is known about the physiological functions of the exonuclease domain [10]. In *Drosophila*, the exonuclease domain of WRN protein is encoded by *WRNexo* (CG7670; FBgn0038608) [10, 12]. The deletion null mutant flies for *WRNexo* removing most of its gene sequences (*WRNexo^Δ^*), are fertile and viable despite some developmental defects [10]. The mutants recapitulate accelerated ageing phenotypes of WS with an increased tumor incidence and shortened lifespan, revealing critical roles of the exonuclease domain in WS [10]. Interestingly, the mutants also display reduced fat contents, abnormal activity/sleep patterns, and reduced resistance to some environmental stresses such as starvation and thermal stress [3, 13]. These observations suggest that the exonuclease domain also plays an important role in metabolism and systemic physiology beyond its primary function in genome stability.

Dietary restriction (DR) or caloric restriction (CR), reduction of food or calorie intake without causing malnutrition, is one of the most robust non-genetic interventions that delays aging and extends the lifespan of a variety of model organisms, including rhesus monkeys and fruit flies [14]. DR also promotes health by mitigating many age-related disease symptoms in both humans and animal models of human disease [15]. As such, DR can be a potential non-genetic treatment option for WS patients. In rats, it was shown that CR increases the WRN protein level in the liver [16], functionally homologous to the fat body in *Drosophila*, suggesting that diet can regulate the expression of WRN. Importantly, the fat body plays critical roles in mediating lifespan extension by DR in flies [17, 18]. Yet, whether DR could ameliorate accelerated aging symptoms and delay premature mortality of WS has not been tested to date. Although the molecular mechanisms of DR are not fully understood, recent research has shown that nutrient sensing pathways play important roles in DR effects [19]. In flies, a number of studies have shown that restriction of dietary yeast, the main protein source in the diet, is sufficient to extend lifespan [20–25], indicating that molecular pathways related to protein sensing and/or metabolism may limit the beneficial effects of DR. This also implies that DR could extend lifespan and delay accelerated aging processes of fly models of WS if their protein sensing mechanisms are not impaired.

Here, we investigate the physiological roles of the exonuclease activity of WRN in DR/diet-mediated lifespan, stress resistances, and sleep/activity patterns using *WRNexo^Δ^* null mutant flies [3, 10, 13]. Our findings show that the *WRNexo^Δ^* mutant flies display a significantly altered physiological response to diet, including a deleterious effect on lifespan by DR in female flies, compared to WT flies. This work contributes to the understanding of the benefits and costs of dietary interventions in WS patients.

## Materials and methods

### Fly Stocks and Maintenance

*WRNexo^Δ^* null mutants (*w^1118^/ w^1118^; +/+; WRNexo^Δ^/TM6, Sb, GFP*) and their isogenic *w^1118^*wildtype control flies were generously provided by Elyse Bolterstein [10, 13]. Flies were expanded in vials (23 mm X 95 mm) containing ∼ 5 mL of a standard cornmeal-based medium for larval growth (Genesee Scientific, Cat#: 66-113) adapted from the Bloomington Drosophila Stock Center (https://bdsc.indiana.edu). All flies for growth/expansion and experiments were maintained at 25°C under a 12 h:12 h light–dark cycle with 40∼60% relative humidity conditions. For all experiments performed, ∼ 48 hour cohorts after eclosion were collected and allowed an additional one day of post-eclosion mating on the cornmeal-based medium before being separated by sex and genotype (homozygous *WRNexo^Δ^* mutants) on light CO_2_ anesthesia. Separated flies were then transferred by sex to respective sucrose-yeast experimental food vials with fixed 5% [w/v] sucrose (Genesee Scientific, Cat#: 62-112) and varying yeast (Brewer’s yeast, MP Biomedicals) concentrations (1%, 5%, 20% [w/v]) [26]. For relative comparison, 1%, 5%, 20% yeast diets were denoted as 1Y malnutrition (Mal) diet, 5Y (DR) diet, and 20Y control (Con) diet, respectively. For yeast supplementation experiment in figure 4, yeast paste was prepared from a 1:1.25 [w/v] yeast (Brewer’s yeast, MP Biomedicals) to water ratio. Experimental food vials were prepared in batches, wrapped with plastic bags to prevent dehydration, and stored in 4°C refrigerators until use. Only fresh food vials less than 2 weeks old were used for all assays.

### Lifespan Assays

Seven to ten replicate vials with up to ∼ 25 flies per replicate vial were set up for each diet, genotype, and sex. Flies were transferred to fresh vials every 2∼3 days (usually 2 days) for figure 1 and every 2 days for figure 4. At each transfer, dead flies were counted and recorded on physical data forms before statistical analysis using JMP 14 (SAS Inc.) statistical package and the online analysis pipeline OASIS2 (https://sbi.postech.ac.kr/oasis2/) [27]. Lost or dead/damaged flies stuck on the cotton ball during transfer were censored from analysis. For figure 4, a half spatula-full of refrigerated yeast paste (less than 1 week old after preparation) was smeared on the side of DR/5Y food vials a few hours before each transfer. To measure accurate mortality pattern in figure 4, transfer was performed at the same time of day (between 8am – 10am).

### Stress Assays

For both starvation and oxidative stress assays, separated flies (∼48 hour cohorts followed by one day of post eclosion mating) were aged for 9 ∼ 10 days in 5Y and 20Y food vials until the day of the assays. Due to an extremely high early mortality particularly in *WRNexo^Δ^* mutants, the 1Y malnutrition diet was not tested for stress response. Flies (6 replicates with 10 ∼ 20 flies in each replicate) of each condition were transferred to starvation vials (1.5% agar (Genesee Scientific, Cat#: 66-103) as water source) and oxidative stress vials with paraquat (Methyl viologen dichloride hydrate; Sigma, Cat#: 856177) solution (10 mM paraquat in 10% [w/v] sucrose solution). For each oxidative stress vial, one piece of Kimwipe (Kimberly-Clark Professional™) soaked with 2 mL of the paraquat solution was placed into the vial, then two round filter papers (23 mm, Whatman) were pushed into the vial until the filter papers were damp with paraquat solution. The excess paraquat solution was wiped off the sides of the vial. To control artifacts caused by potentially different feeding rhythms of flies in different conditions, flies for oxidative stress were first wet-starved for 5 hours in starvation vials (1.5% agar) before transferred to paraquat solution vials. For both stress assays, the number of dead flies in each vial was counted every 2-8 hours until all of the flies were dead.

### Sleep/Activity measurement

Monitoring of locomotor activity and sleep (defined as inactivity for 5 or more minutes) was assayed using the Drosophila Activity Monitoring (DAM, Trikinetics) system following established protocols [28–30]. Briefly, the DAM2 monitors contain 32 channels in which glass tubes containing food and flies are loaded. Each channel records the fly’s movements continuously by shining an infrared beam through the center of the glass tube and recording the number of beam breaks. Flies aged for ∼7 days on each diet were loaded into glass tubes containing corresponding diet after light CO_2_ anesthetization. The activity and sleep of single flies were monitored for a period of ∼ 3 days, resulting in a total duration of diet treatment ∼ 10 days at the end of sleep measurement, which is comparable to the flies used for stress assays. To remove potential effects of CO_2_ anesthesia influencing the sleep and activity of flies, the first experimental day (day 8) was discarded from sleep/activity analysis. Also, flies with less than 10 beam breakings during the last experimental day, which is usually an indication of damaged flies due to moisture condensation in the glass tubes, were also removed from analysis. Activity and sleep profiles and parameters were generated with the Counting Macro program [29].

### Statistics

Descriptive survival statistics (such as mean lifespan and mortality rates) and pair-wise comparison of survival curves between groups (log-rank test) for lifespan and stress resistance assays (Fig. 1, 2, and 4; Supplementary Data 1, 2, and 4) were obtained using OASIS2 (https://sbi.postech.ac.kr/oasis2/) [27] and JMP 14 (SAS Inc.) statistical package. Statistical significance for sleep parameters (Fig. 3) was determined using a two-way ANOVA following the Tukey HSD post-hoc test.

**Figure 1.**
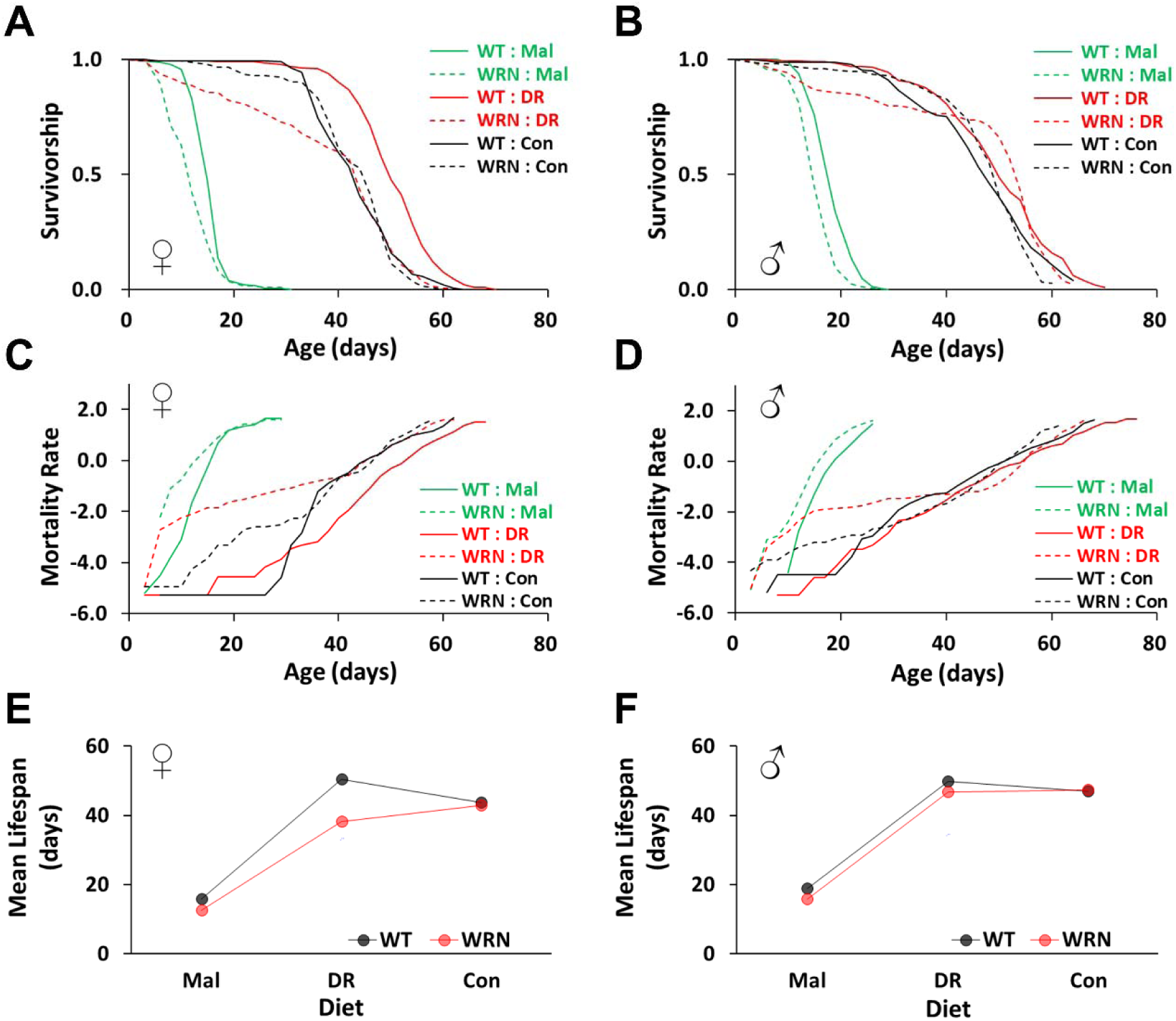
Effects of yeast-restriction diets on the lifespan of Wild-Type (WT) and WRNexo^Δ^ (WRN) mutant flies. (A, B) Survivorship. (C, D) Mortality rate H(t): the risk of death at a given time (t) calculated using the following formula: H(t) = −log(S(t)), where S(t) represents the survival function estimated with the Kaplan-Meier estimator. (E, F) Mean lifespan response to three different yeast-restriction diets. (A, C, E): Males; (B, D, F): Females. Mal (1Y): 1% yeast with 5% sucrose, DR (5Y): 5% yeast with 5% sucrose, Con (20Y): 20% yeast with 5% sucrose. Refer to the main text and Supplementary Data 1 for descriptive statistical analysis and sample sizes.

**Figure 2.**
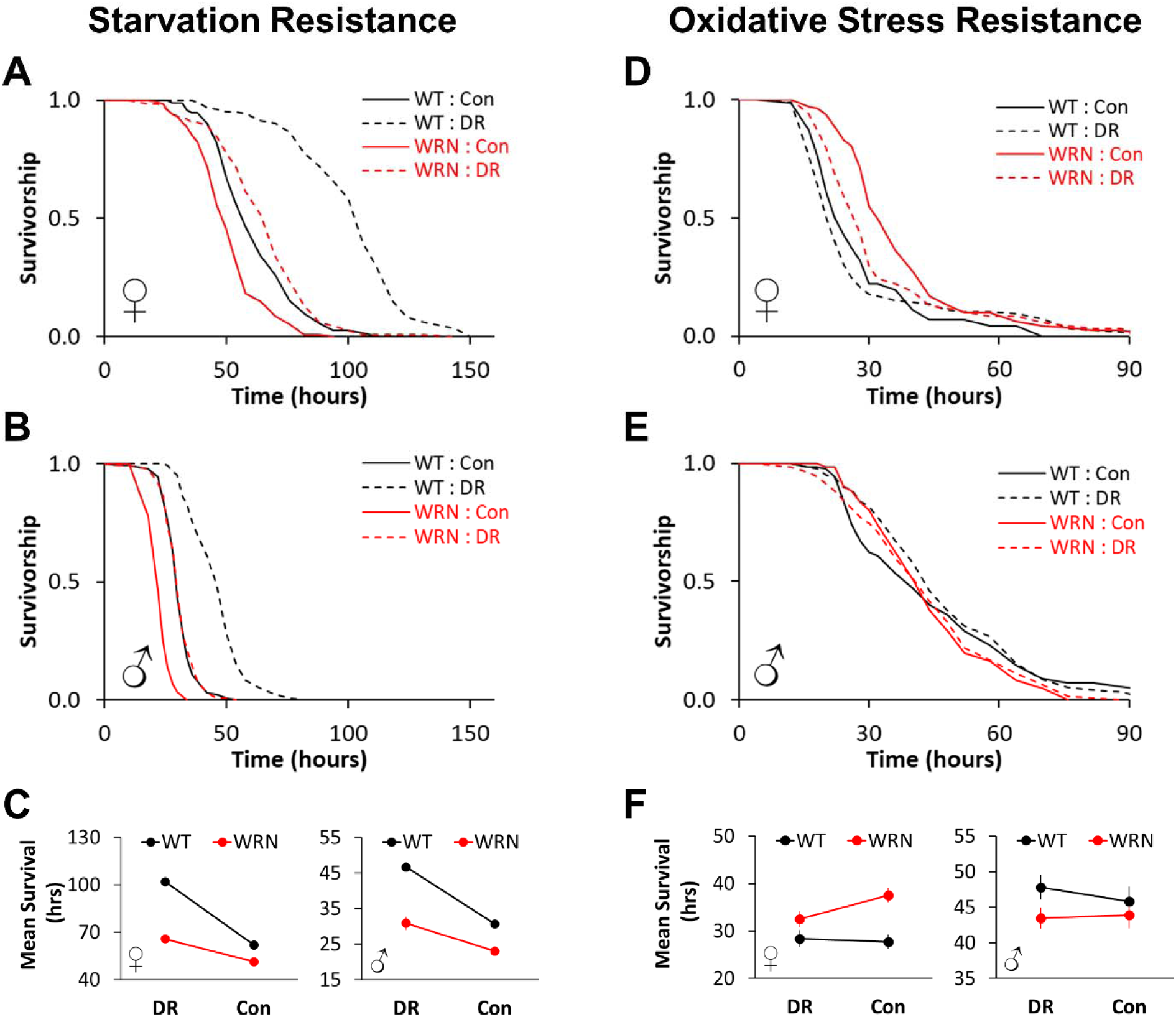
Effects of yeast-restriction diets on the stress response of Wild-Type (WT) and WRNexo^Δ^ (WRN) mutant flies. (A - C) Starvation Resistance. (D, E, F) Oxidative Stress Resistance. (A, D) Survivorship of females. (B, E) Survivorship of males. (C, F) Mean survival changes in response to diets. DR (5Y): 5% yeast with 5% sucrose, Con (20Y): 20% yeast with 5% sucrose. Refer to the main text and Supplementary Data 2 for descriptive statistical analysis and sample sizes.

**Figure 3.**
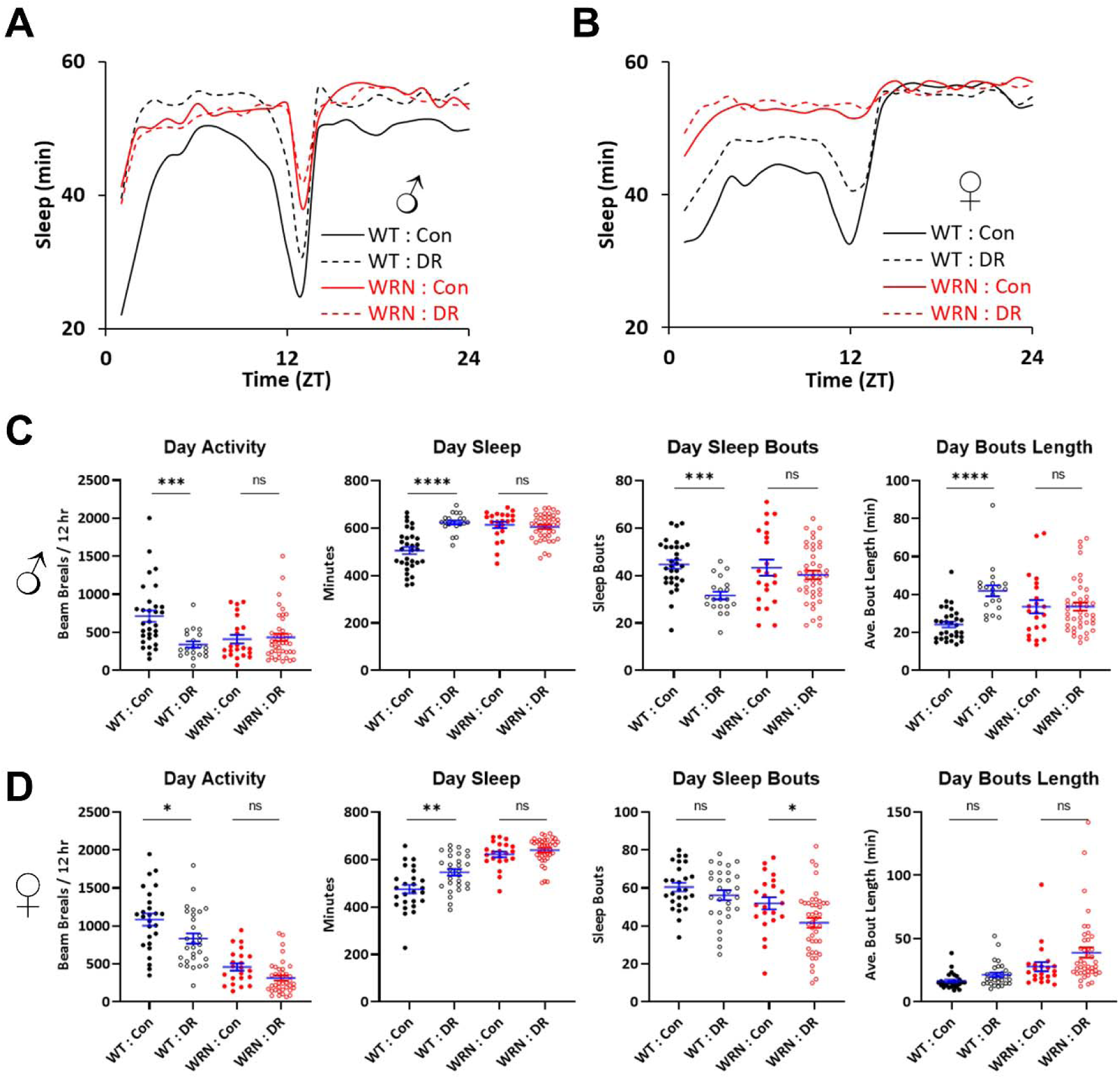
Effects of yeast-restriction diets on the activity and sleep patterns of Wild-Type (WT) and WRNexo^Δ^ (WRN) mutant flies. Flies were fed on DR (5% yeast with 5% sucrose) or Con (20% yeast with 5% sucrose) diets for ∼7 days in vials before sleep measurements were initiated using the DAM system. Sleep was monitored for about 3 days on each diet. ZT0: light on, ZT12: light off. The first day of sleep data was excluded from the analysis due to CO2 anesthesia. (A, B) Twenty-four-hour sleep profiles (average sleep duration per hour plotted in minutes sleep) for male (A) and female (B) flies of WT and WRNexo^Δ^ (WRN) mutants. ZT 0-12 and ZT 12-24 correspond to the 12 hours of the day/light and the night/dark phases, respectively. (C, D) Activity and sleep parameters for male (C) and female (D) flies of WT and WRNexo^Δ^ (WRN) mutants during the light/day phase. Activity: The number of fly beam breaks recorded in the DAM machine. Sleep: The average total sleep time in minutes, calculated as the product of sleep bout numbers and sleep bout length in minutes. Sleep Bouts: The total number of sleep episodes, defined as periods of > 5 minutes of inactivity. Bout Length: The average sleep bout length. Asterisks indicate P values resulting from a 2-way ANOVA followed by the Tukey multiple comparisons post hoc test: * p < 0.05, ** p < 0.01, *** p < 0.001, **** p < 0.0001. Refer to the main text and Supplementary Data 3 for descriptive statistical analysis, sample sizes, and the analysis for the night/dark phase.

**Figure 4.**
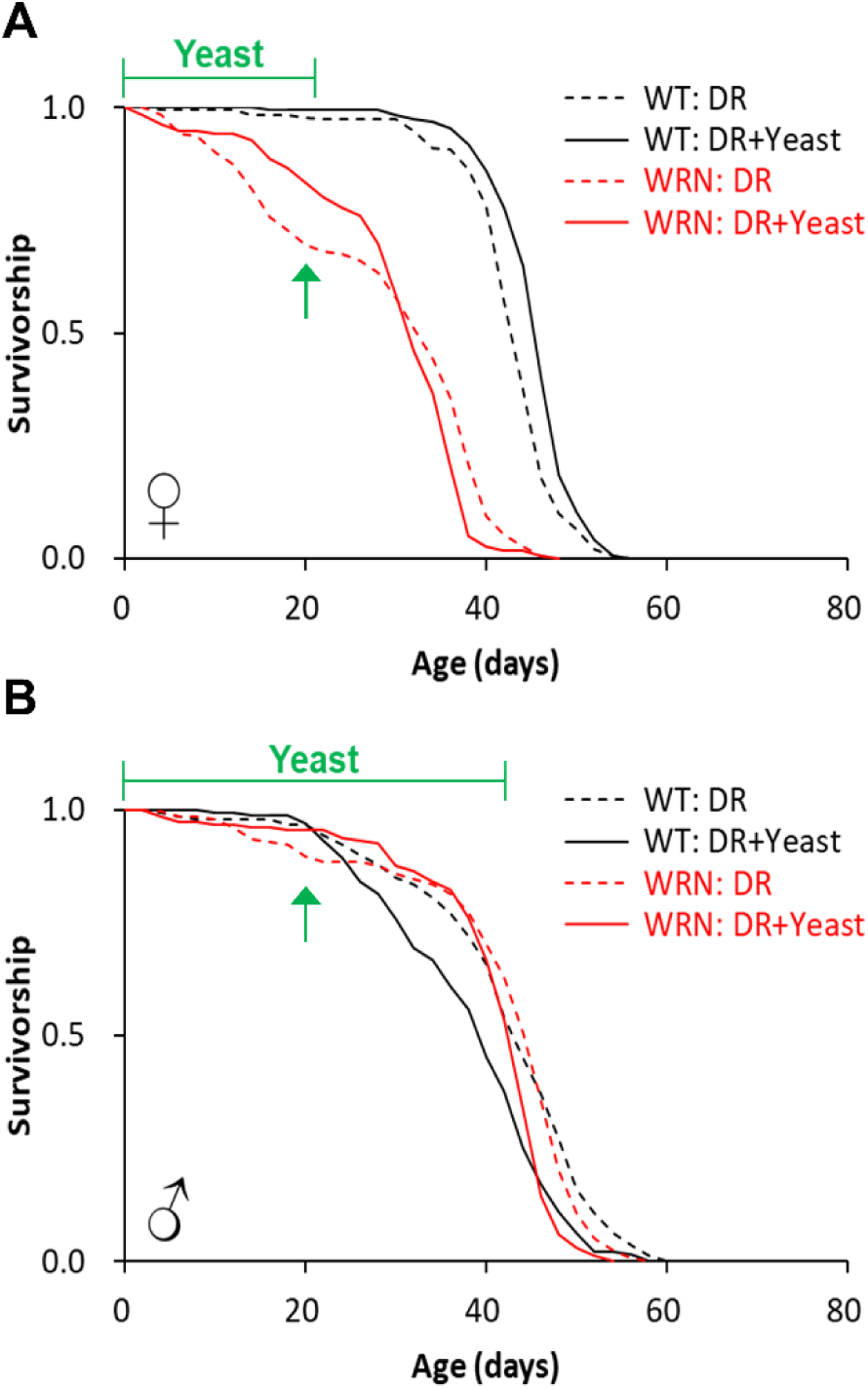
Effects of yeast supplementation on lifespan of Wild-Type (WT) and WRNexo^Δ^ (WRN) mutant flies. Survivorship of female (A) and male (B) flies of WT and WRNexo^Δ^ (WRN) mutants. DR: 5% yeast with 5% sucrose, DR + Yeast: DR diet supplemented with yeast paste (Brewer’s yeast mixed with water in 2.5: 1 ratio). Yeast paste was supplemented fresh on the side of lifespan vials every 2 days when dead flies were counted. Green bars labelled ‘Yeast’ at the top of the graphs represent the duration of time for yeast supplementation. Green arrows indicate representative time points when yeast supplementation increased the survival of the WRNexo^Δ^ (WRN) mutants. Refer to methods and Supplementary Data 4 for descriptive statistical analysis and sample sizes.

## Results

### DR fails to extend lifespan of WRNexo***^Δ^*** mutant flies

A previous study showed that the median lifespan of *WRNexo^Δ^* mutant flies was reduced compared to that of WT flies by 44% in females and 19% in males [3]. To explore how diet impacts the lifespan of *WRNexo^Δ^* mutant flies and also to test whether DR has the potential to delay premature aging and reverse shortened lifespan of the mutants, we used a protein-restriction DR protocol, where yeast (protein source) concentration is diluted while sugar (carbohydrate source) concentration is fixed [24, 26, 31]. Unlike the total dilution protocol where both yeast and sugar are diluted [26], the yeast-dilution DR regimen is known to be devoid of DR-independent mortality caused by desiccation [24] and sugar-induced obesity [32]. Therefore, our diet protocol accurately assesses the impacts of DR on the aging process and other health markers. To understand the range of beneficial and deleterious effects by diets, we employed a reaction norm approach [33] by using three serially diluted diets (1Y/Mal, 5Y/DR, 20Y/Con diets; see materials and methods).

Overall, lifespan and mortality analyses revealed significant gene and diet effects in a sex-dependent manner. In the 1Y/Mal diet, lifespan was dramatically reduced compared to that of 20Y diet in both WT (female: 15.8 days vs 43.7 days; −63.9%, χ2 = 422.3, P < 0.001; male: 18.8 days vs 46.9 days; −59.9%, χ2 = 369.5, P < 0.001) and *WRNexo^Δ^* flies (female: 12.5 days vs 42.9 days; −70.8%, χ2 = 417.7, P < 0.001; male: 15.8 days vs 47.3 days; −66.5%, χ2 = 379.1, P < 0.001), presumably due to malnutrition caused by severe protein restriction in 1Y diet. In the 1Y diet, despite the significant decrease in lifespan of WT flies, the lifespan of *WRNexo^Δ^* flies mutant flies was even shorter than that of WT, indicating that the physiological processes in the mutants are still capable of responding to the diet even under malnutrition conditions (female: 15.8 days (WT) vs 12.5 days (*WRNexo^Δ^*); −20.6%, χ2 = 38.9, P < 0.001; male: 18.8 days (WT) vs 15.8 days (*WRNexo^Δ^*); −15.8%, χ2 = 41.5, P < 0.001) (Fig. 1 and Supplementary data 1).

Surprisingly, in protein-rich 20Y/Con diet, the mean lifespan of *WRNexo^Δ^* mutant flies was not reduced compared to that of WT flies (female: 43.7 days (WT) vs 42.9 days (*WRNexo^Δ^*), −1.7%, χ2 = 0.1, P = 0.705; male: 46.9 days (WT) vs 47.3 days (*WRNexo*), 0.9%, χ2 =0.3, P = 0.589) (Fig. 1 and Supplementary data 1). These results show that the composition of food impacts the lifespan of *WRNexo^Δ^* mutant flies and suggest that macro- and/or micro-nutrients in yeast can slow down or even reverse the accelerated premature death of *WRNexo^Δ^* flies. As expected, DR/5Y diet significantly extended lifespan compared to Con diet in WT (female: 15.5%, χ2 = 62.2, P < 0.001; male: 6.4%, χ2 = 6.8, P = 0.009) (Fig. 1 and Supplementary data 1). DR was more effective in increasing lifespan in females, which is consistent with previous reports that females typically display larger lifespan increases by DR in flies [34, 35]. However, DR failed to extend lifespan of *WRNexo^Δ^* flies in both sexes (Fig. 1 and Supplementary data 1). The mean lifespan of *WRNexo^Δ^* mutants in DR diet was even shorter than that of Con diet in both sexes with little to no statistical significance (female: −11%, χ2 = 0.9, P = 0.335; male: −0.9%, χ2 = 3.2, P = 0.074). Notably, similar to the previous report by Cassidy et al. [3], the overall impact of the WRNexo mutation on mean lifespan was stronger in female mutants. Although male *WRNexo^Δ^* mutants displayed increased early mortality on the DR diet compared to male WT flies, their mean lifespan was comparable to that of WT flies. Furthermore, this increased early mortality of male *WRNexo^Δ^* mutants on the DR diet disappeared on the Con diet, implying that dietary yeast protects against early mortality in the mutants (Fig. 1 and Supplementary data 1). Together, these results indicate that the lifespan responses to dietary yeast is altered in *WRNexo^Δ^* mutant flies, with a sexually dimorphic lifespan pattern. They also suggest that a functional WRNexo protein plays a role in the lifespan responses to dietary yeast and is required for DR-mediated lifespan extension in flies.

### WRNexo***^Δ^*** mutants display altered diet-dependent stress resistance

As DR was incapable of lifespan extension and was even deleterious for the lifespan of *WRNexo^Δ^* flies, leading to a higher rate of early mortality during the first ∼3 weeks (Fig. 1A and 1C), we hypothesized that the physiological and metabolic adaptations linked to lifespan extension by DR may also be altered or impaired in the mutants. To test this hypothesis, we first evaluated starvation resistance of *WRNexo^Δ^* flies along with their control flies after ∼ 10 days on DR and Con diets. The 1Y/Mal diet was not tested due to high mortality in both WT and mutants. Since DR administered with a reduction in dietary yeast, as per the protocol employed in this study, increases starvation resistance in WT flies [25, 36], given their lifespan responses to DR, we predicted that *WRNexo^Δ^* mutant flies would exhibit an altered starvation resistance pattern on these diets. In both sexes, *WRNexo^Δ^* mutants were more sensitive to starvation stress and died faster than WT flies in both DR (female: 102.0 hours (WT) vs 65.6 hours (*WRNexo^Δ^*); −35.7%, χ2 = 136.4, P < 0.001; male: 46.6 hours (WT) vs 30.8 hours (*WRNexo^Δ^*); −33.9%, χ2 = 148.0, P < 0.001) and Con (female: 63.0 hours (WT) vs 51.3 hours (*WRNexo^Δ^*); −35.7%, χ2 = 136.4, P < 0.001; male: 30.8 hours (WT) vs 23.0 hours (*WRNexo^Δ^*); −25.1%, χ2 = 93.8, P < 0.001) diets (Fig. 2A-C and Supplementary data 2). This observation is largely consistent with a recent report [13], although a standard cornmeal-based diet instead of sucrose-yeast (SY) diet was used in the study. As expected, DR significantly increased starvation resistance in WT female and male flies by 64.6% (mean survival: 62.0 hours (Con) vs 102.0 hours (DR), χ2 = 138.5, P < 0.001) and 51.6% (mean survival: 30.8 hours (Con) vs 46.6 hours (DR), χ2 = 149.6, P < 0.001), respectively (Fig. 2). In *WRNexo^Δ^* flies, however, DR resulted in a smaller increase in starvation resistance compared to that of WT. The increase of starvation resistance by DR was only 27.9% (mean survival: 51.3 hours (Con) vs 65.6 (DR) hours, χ2 = 44.3, P < 0.001) and 30.8% (mean survival: 23.0 hours (Con) vs 30.8 (DR) hours, χ2 = 30.9, P < 0.001) in female and male mutants, respectively. This pattern corresponds to the lifespan results in figure 1 and suggests that a functional WRNexo protein is required for the full benefit of DR to increase resistance to starvation stress.

Next, using the same rearing conditions as those used for assessing starvation resistance, we measured oxidative stress resistance. Despite shortened lifespan and reduced resistance to starvation stress, a previous study showed that *WRNexo^Δ^* females are more resistant to oxidative stress (5% H_2_O_2_) partially due to increased activities of antioxidant defense system [13]. Given the observation that the effects of DR on lifespan and starvation resistance were less pronounced in *WRNexo^Δ^* flies than WT flies, we hypothesized that *WRNexo^Δ^* flies might display greater sensitivity to oxidative stress when reared in DR diet compared to Con diet. To test this hypothesis, we chose paraquat (methyl viologen; 10 mM in 10% sucrose solution; see materials and method) to induce oxidative stress [13] as our preliminary RNA-Seq analysis discovered that multiple cytochrome P450 genes, known to be involved in detoxifying exogenous chemicals, were strongly mis-regulated in *WRNexo^Δ^* flies (data not shown). In males, neither genotype (WT vs *WRNexo^Δ^*) nor diet (Con vs DR) showed significant differences in survival after paraquat treatment (Fig. 2 D-F and Supplementary data 2). However, similar to the report by Epiney et al [13], female *WRNexo^Δ^* flies were more resistant to paraquat in both Con (27.7 hours (WT) vs 37.5 hours (*WRNexo^Δ^*); 35.4%, χ2 = 21.2, P < 0.001) and DR diets (28.4 hours (WT) vs 32.5 hours (*WRNexo^Δ^*); 14.7%, χ2 = 8.7, P = 0.003) (Fig. 2D-F and Supplementary data 2). Notably, while mean survival of WT flies on paraquat was not significantly different between Con and DR (28.4 hours (Con) vs 27.7 hours (DR); χ2 = 0.1, P = 0.737), that of *WRNexo^Δ^* flies was lower in DR than in Con diet by ∼ 13% (37.5 hours (Con) vs 32.5 hours (DR); χ2 = 7.2, P = 0.007). This suggests that, unlike WT flies where the yeast concentration difference between DR and Con diets (5% vs 20%) does not lead to varying resistance to oxidative stress, lower yeast concentration in the DR diet reduces resistance in *WRNexo^Δ^* mutant flies. Therefore, similar to lifespan and starvation resistance results, the oxidative stress resistance pattern strongly indicates that *WRNexo^Δ^* flies have impaired physiological responses to diets.

### WRNexo***^Δ^*** mutants display altered diet-dependent sleep repatterning

In flies, the absence or quiescence of locomotive activity beyond a given time period (5 minutes), is considered sleep or rest [29, 37]. This approach displays several key analogous features of mammalian sleep, including an increased arousal threshold, regulation by circadian clocks, and homeostatic needs [38]. It has been successfully used to dissect the genetic and neuronal mechanisms of sleep. Similar to mammals, environmental changes such as in nutritional composition and value in the food affect locomotor activity and sleep patterns in flies [37, 39, 40]. Since *WRNexo^Δ^* mutant flies displayed altered lifespan and stress resistance responses to diets (Fig. 1 and 2), we hypothesized that the mutants’ activity/sleep patterns in response to yeast dilution is also disrupted. After pre-treating the flies on DR and Con diets for ∼ 7 days, we monitored time-dependent locomotor activity and sleep patterns of the flies on DR and Con diets for ∼ 2 days. We analyzed the total activity level (the number of flies’ beam breaks in the DAM machine) and total sleep time (the product of sleep bout numbers and sleep bout length in minutes) separately for daytime and nighttime.

Compared to Con diet, DR diet significantly reduced the total activity of WT flies in both sexes (males: 713.8 (Con) vs 340.7 (DR), P < 0.001, females: 1086 (Con) vs 835.1 (DR), P = 0.0165), particularly during the daytime, which led to increased total sleep during the daytime (males: 505.2 minutes (Con) vs 623.6 minutes (DR), P < 0.0001, females: 475.1 minutes (Con) vs 546.3 minutes (DR), P = 0.0013) (Fig. 3 and Supplementary data 3). This DR-dependent increase in daytime sleep in WT flies was primarily due to decreased sleep bout numbers (males: 44.7 (Con) vs 31.6 (DR), P < 0.001) with increased sleep bout lengths (males: 24.2 minutes (Con) vs 42.0 minutes (DR), P < 0.0001) particularly in males (Fig. 3C and Supplementary data 3). In females, although the trend of change in sleep bout numbers and sleep bout length due to DR was similar to that in males, the differences between the Con and DR diets were not statistically significant (Fig. 3D and Supplementary data 3).

As predicted, unlike WT flies, the re-patterning of activity and sleep by dietary yeast was strongly suppressed in *WRNexo^Δ^* mutants. There were negligible differences between Con and DR diets in total activity (males: 411.1 (Con) vs 434.8 (DR), P = 0.992, females: 461.4 (Con) vs 316.5 (DR), P = 0.2944) and total sleep (males: 613.5 minutes (Con) vs 605.0 minutes (DR), P = 0.9565, females: 621.4 minutes (Con) vs 639.6 minutes (DR), P = 0.7567) during the daytime. In male mutants, unlike WT flies, neither sleep bout numbers nor sleep bout lengths were affected by DR (Fig. 3 and Supplementary data 3). These observations indicate that *WRNexo^Δ^* mutant flies have lost the mechanisms to regulate activity and sleep in response to dietary yeast. Thus, these results also suggest that, in addition to altered baseline sleep in *WRNexo^Δ^* mutants as observed previously [3, 13], a functional exonuclease activity of WRN protein is required for diet-mediated sleep regulation.

### Yeast supplementation delays accelerated mortality of WRNexo***^Δ^*** mutants

Above concentrations that cause malnutrition, dietary yeast limits lifespan; the higher the yeast concentration, the shorter the lifespan [26, 36, 41]. Since DR did not extend lifespan of *WRNexo^Δ^* mutant flies, resulting in a significant increase in early mortality, we hypothesized that *WRNexo^Δ^* mutant flies require higher nutritional intake for survival during their early stages in adult compared to WT flies. To test this possibility, we examined the effect of additional dietary yeast on the lifespan of *WRNexo^Δ^* mutants. Flies were supplemented with a concentrated yeast paste (see Materials and Methods) to DR diet for 3 weeks for females and 6 weeks for males (Fig. 4). The yeast paste was supplied on the side of food vial such that flies can consume both yeast paste and DR diet *ad libitum*. As predicted, yeast supplementation delayed and was protective against the early mortality observed in *WRNexo^Δ^* flies on the DR diet (Fig. 4; indicated by arrows). For example, at day 20, yeast supplementation increased survival of mutants on DR diet by 14% and 6% in females and males, respectively, while it had no obvious impacts on survival in WT flies. Similarly, the day of 25% mortality was delayed from 18 day (DR) to 28 day (yeast supplementation) in female *WRNexo^Δ^* flies. In male *WRNexo^Δ^* flies, the day of 25% mortality was 40 day for both groups, which is comparable to that of WT on DR diet (38 day). Overall, these data indicate that, despite the overall pro-aging effects of yeast in WT flies [25, 26, 42], macro- and/or micronutrients in yeast during early life can at least partially reverse the accelerated aging process of *WRNexo^Δ^* flies. Given that protein is a major component of the dietary yeast in the yeast paste, our findings also raise the possibility that supplementing protein and/or specific amino acids might be able to delay the accelerated aging in *WRNexo^Δ^* flies.

## Discussion

WS is caused by mutations in the human WRN gene, which contains two important functional domains, a RecQ-type helicase domain and an exonuclease domain. At present, there is no cure for WS, which highlights dietary and lifestyle interventions, such as DR, as potential options for increasing the life expectancy of individuals with WS. In this study, utilizing a deletion null mutant *Drosophila* (*WRNexo^Δ^*) for the *WRNexo* gene encoding a protein homologous to the exonuclease domain of the human WRN gene [3, 13], we demonstrate that a yeast-restriction DR regimen fails to extend the lifespan of a fly model of WS. We also show that *WRNexo^Δ^* mutants display significantly altered overall physiological (stress resistance) and behavioral (activity/sleep) responses to DR compared to those of WT control flies (Fig. 1-3). Therefore, our observations provide direct evidence that the WRNexo protein is necessary for a wide range of DR responses in WT flies. Together with recent studies showing its role in lipid metabolism [3, 13], our data also raise the possibility that the exonuclease domain of the WRN protein may be directly involved in nutrient sensing and metabolism, particularly amino acids given the type of DR regimen used in our study. However, our results may also indicate that DR affects the WRN protein’s primary functions in genome stability and DNA-related molecular processes, including replication, repair, recombination, and transcription, thereby indirectly altering the DR response in lifespan and physiology observed in *WRNexo^Δ^* mutants. While it is only speculative, we predict that this possibility is unlikely, as it was shown that lifespan extension by DR is not attributed to protection against DNA damage in flies [43] and is still able to delay accelerated aging in DNA repair-deficient mice [44].

Notably, the mean lifespan of female *WRNexo^Δ^* mutants on the DR diet (5Y) was even shorter than that of the protein-rich Con diet (20Y), primarily due to increased early mortality during the first ∼ 2 weeks (Fig. 1A, C, and E). This observation indicates that nutrient-restricted diets such as the 5Y diet used in this study, which are beneficial in WT flies, can have deleterious effects in the fly model of WS. It also implies that *WRNexo^Δ^* mutant flies require more nutrients than the amount that confers lifespan extension in WT flies. In accordance with this, supplementation of protein-rich yeast pastes partially rescued the high early-life mortality (Fig. 4). This finding provides additional evidence that certain nutrients present in dietary yeast can effectively decrease mortality in *WRNexo^Δ^* mutants. Considering that dietary yeast in our fly food serves as the primary source for protein [26, 37], which is known to limit lifespan and also play as the major dietary component in DR-mediated lifespan modulation in flies [25, 26, 42], we favor the idea that the molecular pathways involved in protein metabolism and sensing might be compromised in *WRNexo^Δ^* mutant flies. However, it’s also possible that the residual amounts of other macro- and micronutrients in dietary yeast, such as lipids and vitamins, could have contributed to the phenotypes of the mutants [45, 46]. For example, previous studies demonstrated that vitamin C and NAD^+^ could at least partially rescue the shortened lifespan of WRN mutants in some model organisms [5, 47]. A further study employing a defined medium that controls the concentrations of total protein and each amino acid, along with vitamin C or NAD^+^ supplementation, could provide a clearer answer to this possibility.

Elevated resistance against environmental stress is often associated with increased lifespan. While age functions as a confounding variable impacting stress resistance [48], the composition of diets, including yeast and sugar concentration, significantly affect the stress resistance of flies [25, 48–50]. For example, yeast-restriction DR increases survival during starvation stress, partly due to elevated lipid content (synthesized via lipogenesis) and enhanced utilization (broken down through lipolysis) [24]. Earlier studies indicates that *WRNexo^Δ^* mutants contain a significantly lower amount of body fat and are more sensitive to starvation stress compared to those of WT flies in both sexes. [3, 13]. However, only female mutants unexpectedly exhibit increased resistance to oxidative stress, attributed to the upregulation of cellular antioxidant defense mechanisms [3, 13]. Our observation is largely consistent with these findings: Regardless of diets, *WRNexo^Δ^* mutants exhibited greater sensitivity to starvation stress in both sexes, while displaying increased resistance to oxidative stress only in females compared to WT control flies (Fig. 2). With respect to the diet effects, although DR was still able to enhance the starvation resistance of *WRNexo^Δ^* mutant flies, the increase was much smaller than that was observed in WT flies in both sexes (Fig. 2A and 2B). Unlike starvation stress resistance, diet has minimal impact on oxidative stress resistance in both sexes of WT flies and in males of the mutants.

Interestingly, DR negatively affects oxidative stress resistance in female mutants. This finding is consistent with an earlier study [48] and also comparable to their lifespan on DR and Con diets (Fig. 1). Taken together, although stress-specific with sexually dimorphic patterns, our data indicates that WRNexo plays a role in diet-dependent stress response. In line with these data, it is worth noting that our preliminary RNA-Seq transcriptome analysis identified that multiple pathways involved in lipid metabolism and detoxification of exogenic stressors were differentially expressed in *WRNexo^Δ^* mutant flies in a diet-dependent manner (data not shown), which provide molecular signatures that may explain the phenotypes of the mutants reported in this study. Also, considering the lifespan patterns of the mutants between DR and Con diets and compared to those of WT flies, these observations imply that the mutants’ altered response to diets in starvation and oxidative stress resistance cannot fully explain their lifespan patterns.

There are some discrepant observations regarding the impacts of dietary sugar and yeast in fly food for sleep architecture [37, 39, 40]. Our data appears to align with one of the earlier studies [39], revealing that dietary yeast tends to fragment sleep by increasing bout numbers while decreasing bout length, particular in male WT flies (Fig. 3C and 3D). Although it is still not well understood whether lowered sleep quality caused by a protein-rich diet has a negative impact on lifespan or if yeast-driven repatterning of sleep contributes to DR-mediated lifespan modulation, our data indicate that the mechanisms by which flies sense the nutritional values of yeast to regulate sleep is strongly suppressed in *WRNexo^Δ^* mutant flies. It is worth noting that, unlike the lifespan patterns, the altered responses to dietary yeast in activity and sleep patterns in WT flies were more evident in males than in females. This raises the possibility that the repatterning of activity and sleep by DR is unlikely the key determinant of lifespan extension through DR.

Overall, using *WRNexo^Δ^* mutant flies, our data suggest previously uncharacterized roles of the exonuclease domain of the human WRN protein in response to diets to regulate lifespan, and stress response, and activity/sleep. Considering that the *Drosophila* WRNexo protein shares many molecular and biochemical characteristics of the human WRN protein [3, 12, 13, 51–53], we suggest that nutritional interventions and treatments for individuals with WS should be performed with care. Further studies aiming to identify molecular signatures, such as differentially regulated genes and metabolites, between WT and *WRNexo^Δ^* mutant flies on varied diets, investigating whether compromised genome stability in the mutants is influenced by diets, and determining the specific nutrients that impact the mutants’ lifespan, hold significant importance for translational insights for WS patients.

## Acknowledgements

We thank the Elyse Bolterstein lab for providing the mutant flies, and we also appreciate the technical assistance of Sarayu Alli, Nicholas Wright, Jason Ho, and Erica Hassoun in conducting lifespan and sleep assays. Additionally, we acknowledge the valuable feedback provided by Rafael Demarco, Lee Thompson, and Cynthia Corbitt on an earlier version of this manuscript.

Part of this work was performed with assistance of the University of Louisville Genomics Facility, which is supported by NIH P20GM103436 (KY-IDeA Networks of Biomedical Research Excellence), the J. G. Brown Cancer Center, and user fees.

## Author Contributions

Conceptualization: ES and DH, Investigation: ES, RC, BB, MO, and DH, Data curation: ES, BB, and DH, Analysis: ES, BB, and DH, Manuscript preparation: ES and DH, Supervision, funding acquisition, and project administration: DH. All authors have read and agreed to the published version of the manuscript.

## Data Availability Statement

The datasets used in this study are accessible upon reasonable request from the corresponding author.

